# Dynamic intraglomerular neuronal ensembles represent odor identity and concentration

**DOI:** 10.1101/2023.10.05.560824

**Authors:** Mathieu Loizeau, Alexandra Angelova, Harold Cremer, Jean-Claude Platel

**Affiliations:** Aix Marseille University, CNRS, IBDM, UMR 7288, Turing Centre for Living systems, Marseille, France; Aix-Marseille University, INSERM, Inmed, UMR 1249, Turing Centre for Living systems, Marseille, France

## Abstract

The olfactory system has been extensively studied from an anatomical and functional point of view, yet how and where certain basic odor characteristics such as identity and concentration are represented in the brain remains poorly understood. The glomerular layer of the olfactory bulb is the first brain region that integrates olfactory signals and enables a topographic representation of odor identity. We investigated this odor encoding at the intraglomerular network level using genetic labeling, in vivo imaging, and computational methods. Our analyses demonstrated that glomerular glutamatergic neurons encode both odor identity and concentration. Furthermore, in vivo structural and functional imaging of sister neurons revealed the emergence of intraglomerular neuronal ensembles governed by odor identity and concentration. These findings revealed a novel network mechanism that enables the simultaneous coding of odor identity and concentration within glomerular modules, suggesting a potential mechanism for odor decorrelation and a new model for odor information processing in the olfactory bulb.

**One-sentence summary:** Unraveling the scent puzzle: Discovery of intraglomerular neuronal ensembles sheds light on odor representation in the brain.

## Main Text

Olfaction is crucial for animals, yet the precise mechanisms by which the brain processes and distinguishes odor identity and concentration remain elusive (*1*). In the olfactory system, odorant molecules are detected by olfactory sensory neurons (OSNs) and transmitted to the glomeruli in the olfactory bulb (OB) to form topographic odor representations (*2, 3, 4, 5*). However, at the level of the cell bodies of output neurons, such as mitral and tufted cells, the neuronal activity in response to olfactory stimulation becomes decorrelated, meaning that the activity of these neurons is becoming less similar. Some studies have proposed that mitral cell ensembles represent both odor identity and concentration (*6-8*), with some cells coding for identity and others for concentration (*9*). However, this theory has shortcomings because this separate representation of identity and concentration can be completely disrupted between different odors, causing some cells to switch their coding roles. Recent studies using demixed principal component analysis (PCA) have proposed that mitral cells primarily encode odor identity, whereas tufted cells could encode both odor identity and concentration (*10*). Additionally, while olfactory information in the glomerular layer is initially represented by a pattern of activated and inhibited glomeruli, recent studies have uncovered a more intricate intraglomerular coding system implying sister neurons, i.e. neurons connected to the same glomerulus, whose mechanism and role remain elusive (*11, 12, 13*).The glomerular layer is populated by various interneuron subtypes. They are mainly GABAergic, but a small group of glutamatergic interneurons, labeled through the ND6 promoter, project locally in the OB (*14*) and could provide a unique opportunity to better understand this glomerular network.

This study aimed to investigate the representation of odor identity and concentration at the intraglomerular cellular level using a combination of genetic, longitudinal in vivo imaging, and computational methods. By employing dimensionality reduction techniques, we uncovered that glomerular ND6+ neurons encode both odor identity and concentration. We showed that at the intraglomerular level, ensembles of sister ND6+ neurons (sis-ND6 neurons) were progressively recruited by increasing odor concentrations. These neuronal ensembles remained stable over weeks. Moreover, the activation of the same glomerulus by a different odor recruited different ensembles of sis-ND6 neurons. These results suggest the presence of concentration- and odor-dependent intraglomerular neuronal ensembles in the OB, which potentially represent the first mechanism of odor decorrelation. We propose that this novel network mechanism can enable the coding of odor identity and concentration within a glomerular module.

### ND6+ glomerular population encodes odor concentration and identity

To study odor representation in glomerular glutamatergic neurons, we used the transgenic mouse line NeuroD6-Cre/tdTomato/GCaMP6s (Fig. 1A). The ND6 promoter drives Cre expression in all glutamatergic neurons in the OB, including the population of vesicular glutamate transporter positive glomerular neurons (Fig. 1B) (*14*). We used two-photon calcium imaging in anesthetized animals and recorded consecutively odor-evoked responses in three to six imaging planes in the glomerular layer to sample a larger population of ND6+ neurons (Fig. 1C; 191 neurons/animal on average). Olfactory stimulation with individual odors (see Methods) at 0.5% triggered a response in 58 % of the ND6+ neurons (Fig. 1DE, n=12 animals, 2485 cells). These individual responses were highly reproducible in each successive stimulation under our conditions (fig. S1, ANOVA: p=0.97). We observed diverse response profiles in the ND6+ neurons, including activations, inhibitions, and more complex responses (92.9%, 6.9%, and 0.2% respectively, Fig. 1D-F). These response profiles are similar to those described for the apical dendrites of mitral cells recorded at the glomerular level (*2*).

**Fig. 1.**
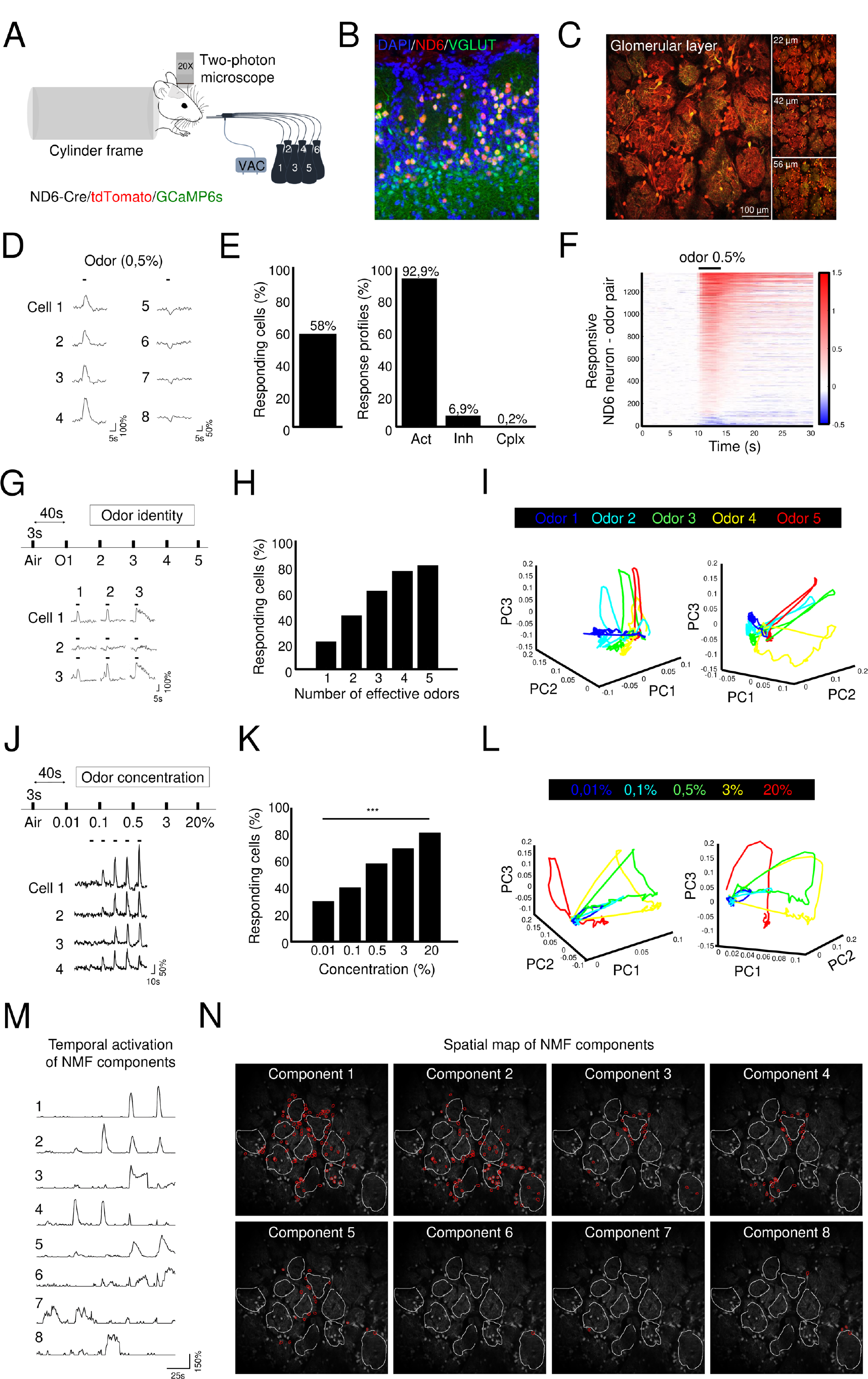
Odor identity and concentration representations in ND6 positive glomerular neurons. (A) Illustration of the odor delivery system and the two-photon imaging configuration in a ND6-Cre/tdTomato/GCaMP6s mouse. (B) Confocal image of a VGLUT2 immunostaining of the glomerular layer in an OB slice from ND6-Cre/tdTomato mice showing extensive colabeling. (C) Representative field of view showing ND6+ neurons in the glomerular layer, captured at 3 different planes with in vivo two-photon imaging. (D) Calcium odor responses observed in ND6+ neurons during 0.5% odor stimulation showing both activation and inhibition responses. (E) (Left) Quantification of the number of responding ND6+ neurons to 0.5% olfactory stimulation (n=12 animals, 2485 ND6+ neurons). (Right) Quantification of response patterns in responding ND6+ neurons, categorized as activations (Act), inhibitions (Inh), or complex responses (Cplx). (F) Raster plot illustrating calcium odor responses in ND6+ neurons during 0.5% odor stimulation (n=12 animals, 1443 ND6+ neurons). (G) Up: experimental protocol of the “odor identity” dataset. Down: calcium activity traces in ND6+ neurons in response to three different odors. (H) Cumulative frequency plot depicting odor tuning, indicating the number of odors eliciting responses out of five tested odors, for 389 ND6+ neurons. (I) Trajectories over time of the ND6+ neuronal population vectors for five different odors, visualized in the space of the first three principal components (PCs). (J) Experimental protocol of the “odor concentration” dataset. Calcium activity traces in ND6+ neurons across a range of odor concentrations. (K) Quantification of the number of responding ND6+ neurons across the concentration range from 0.01 to 20% (n=12 animals, 2485 ND6+ neurons; Cochran-Armitage trend test: p<0.001). (L) Trajectories of the ND6+ neuronal population vectors (from odor stimulation to 45 seconds post-stimulation) for five different concentrations of the same odor, projected onto the first three PCs. (M) Temporal activation profiles of the NNMF components, representing the dynamic patterns of neural activity over concentrations. (N) Spatial map illustrating the distribution of thresholded vectors for each NNMF component, which capture spatial patterns of neural activation within the glomerular layer.

We then investigated whether odor identity could be encoded by this population. We created an “odor identity dataset” that consists of consecutive stimulations with five odors at 1% (see Methods, Fig. 1G; n=5 animals, 389 neurons). For each olfactory stimulation, we observed different populations of responding neurons and forms of responses (Fig. 1G-H). To determine whether odor identity was represented in the ND6+ neurons, we used the classical dimensionality reduction technique known as PCA. The three-dimensional graphical representations of PC1, PC2 and PC3 for each of the five different odors unambiguously show that each odor spanned in a distinct subspace (Fig. 1I and fig. S2), indicating that odor identity was encoded in this population in the glomerular layer.

We next assessed whether the ND6+ neurons also encode odor concentrations. We designed a “concentration range dataset” comprising five consecutive stimulations of a single odor, ranging from 0.01% to 20% (Fig1. J, n=12 animals, 171 glomeruli, 2485 neurons). We observed the individual neuron responses at different concentrations (Fig. 1J). The percentage of responding cells increased significantly from 30% at 0,01% to 81% at 20% (Fig. 1K, n=12 animals, 2485 neurons, Cochran-Armitage trend test: p<0.001). Similarly, this was reflected by a clear increase in the amplitude of the neuronal responses with increasing concentrations (Fig. 1J). To determine whether odor concentration was encoded in the ND6+ neuron population, we performed a PCA. We generated three-dimensional plots of PC1, PC2, and PC3 for five animals (Fig. 1L and fig. S2). In the plot for a single animal, the trajectories associated with different concentrations are situated within distinct subspaces. (Fig. 1L). This observation suggests that the ND6+ neuron population distinctively encodes odor concentration, as indicated by the pronounced separation of trajectories. To extract meaningful patterns of neural activity, we utilized non-negative matrix factorization (NNMF) as an alternative dimensionality reduction technique. We applied NNMF to reveal insightful and informative structures underlying the recorded neural activity. We analyzed the same animal as that shown in Fig. 1L and conducted the NNMF decomposition (see Methods). We plotted the temporal activation of the NNMF components (Fig. 1M) and neurons (on the spatial map) that had the highest weight for each component (see Methods, Fig. 1N). This analysis revealed much more complex cellular behaviors than those observed in the analysis of the percentage of cells recruited or the increase in the amplitude with the concentration. For example, we could distinguish between groups of neurons with responses to higher or lower concentrations (Fig. 1M). Furthermore, we observed intriguing responses, particularly regarding the fourth component in this example, noting a decrease in the magnitude of the neural responses as the odor concentration increased (Fig. 1M). On the spatial map, we have represented the borders of the glomeruli and neurons with the highest weights (see Methods). We observed groups of neurons surrounding several glomeruli in the first component, a group of neurons surrounding one specific glomerulus in the third component, and spatially sparse neurons in other components (Fig. 1N). Although we expected a clear segregation on the basis of only the glomerular map, this repartition seemed to be more complex and pushed us to look at the intraglomerular network.

Overall, we showed that ND6+ neurons in the GL can effectively respond to odor stimulation *in vivo* with different profiles and that they encode both odor identity and concentration. By applying NNMF to temporal and spatial activation analyses, we can unveil more intricate and complex cellular behaviors within their activity patterns.

### *in vivo* morpho-functional characterization of ND6+ sister neurons

On the basis of the results of the NNMF analysis and previous studies(*13*), we investigated the intraglomerular network of the sis-ND6 neurons. The high density of the ND6+-labeled dendrites allowed us to clearly identify the shape of the glomeruli (20 on average in a 600 μm2 field of view, Fig. 2A). By recording high-resolution Z-stack images (Movies S1-2), we distinguished the cell bodies and processes of the ND6+ neurons surrounding each glomerulus (Fig. 2A-B). Benefiting from the superficial location of the glomerular layer, pronounced expression of tdTomato and high quality of our cranial window, we confirmed with high-resolution *in vivo* two-photon imaging that 84% of the labeled cell bodies around the glomerulus had only one main dendrite projecting onto a single glomerulus (Fig. 2B and fig. S3). By tracing this main dendrite, we determined onto which glomerulus each ND6+ neuron was projecting, as illustrated by the colormap (Fig. 2A-B, 21 ND6+ neurons/glomerulus on average over the whole glomerular layer).

**Fig. 2.**
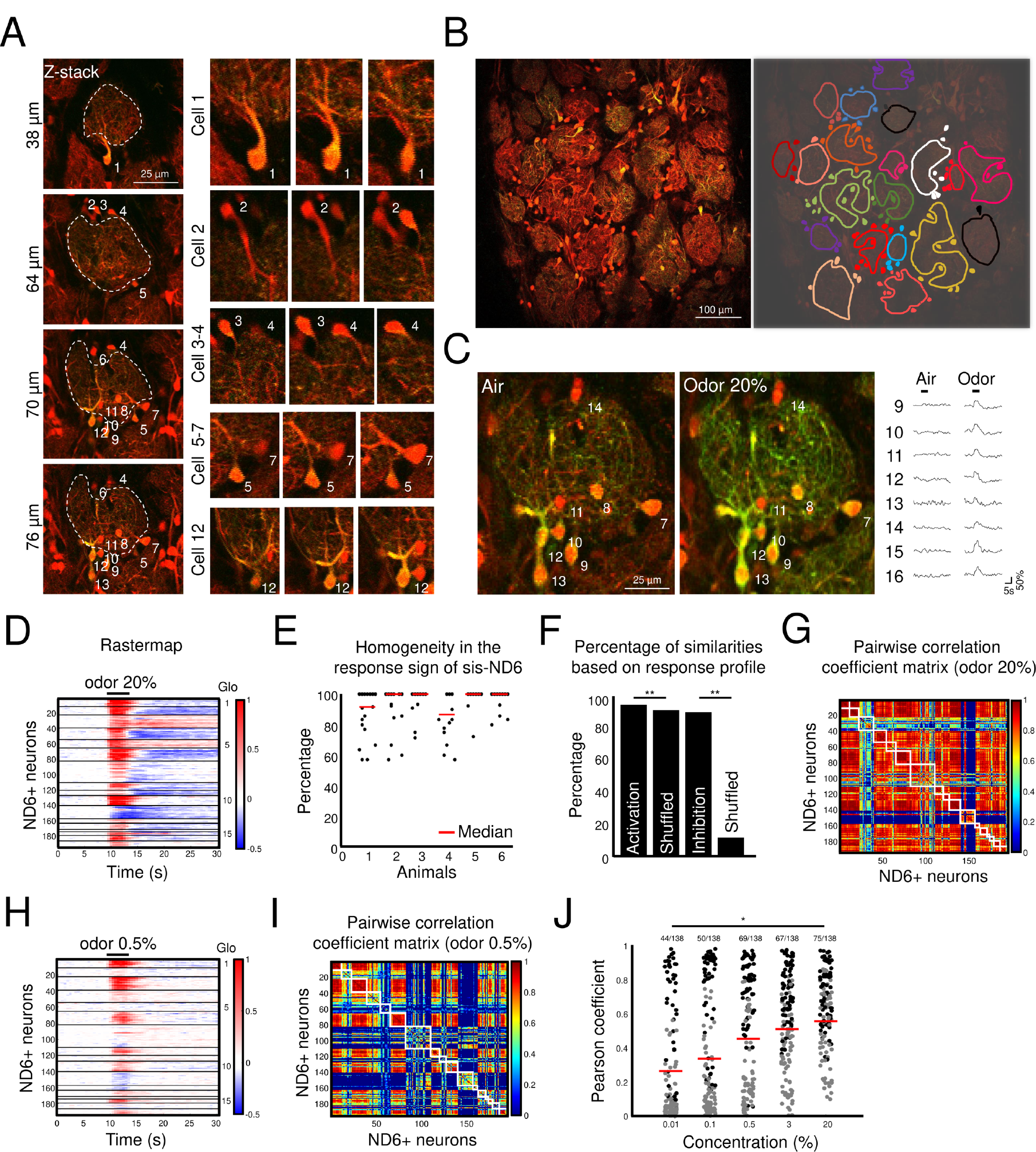
In vivo morpho-functional characterization of sis-ND6 neurons. (A) Close-up view of a specific glomerulus at different depths, illustrating the cell bodies and primary dendrites of sis-ND6 neurons. (B) Reconstruction of intraglomerular ND6 neurons, revealing the spatial arrangement of sis-ND6 neurons within a glomerulus. (C) Zoomed-in view of sis-ND6 neurons before and during olfactory stimulation. The polarity and time course of individual neuron response are displayed on the right. (D) Raster plot depicting calcium odor responses in sis-ND6 neurons from one animal. The black lines delineate the boundaries between glomeruli. (E) Scatter plot illustrating the percentage of similar responses among sis-ND6 neurons within each glomerulus, calculated with respect to the sign of the neuron response (activation, inhibition, complex). The red bar represents the median value (n=6 animals and 127 glomeruli). (F) Bar graph displaying the average percentage of similar responses among sis-ND6 neurons, categorized by the global response of the glomerulus (activation or inhibition; n=6 animals, 127 glomeruli). (G) Pairwise correlation matrix based on the responses of sis-ND6 neurons, highlighting a very high level of correlation between intraglomerular neurons. Each white square corresponds to an intraglomerular ensemble of sis-ND6. (H) Similar to (D), but for 0.5% odor concentration, demonstrating a significant degree of similarity in responses between sis-ND6 neurons. (I) Similar to (G), but for 0.5% odor concentration, indicating that intraglomerular coupling remains robust even at this lower concentration. (J) Mean pairwise correlation of responses of sis-ND6 neurons to olfactory stimulation within a single glomerulus at various concentrations. Statistical tests are made against pairwise correlation of shuffled neurons (10000 iterations). Black and gray dots represent statistically and non-statistically significant mean pairwise correlation. A Chi-squared test shows that the number of significant sis-ND6 increases with concentration (** p<0.05). (n=12 animals, 138 glomeruli).

On the basis of this projection information, we reanalyzed our odor concentration dataset (n=12 animals, 171 glomeruli, 2485 neurons). We sorted the neurons on the raster plot of odor-evoked responses using the projection information of ND6+ neurons. The stimulus-triggered average response of ND6+ neurons was strongly dependent on which glomerulus they projected to, for both 0.5% and 20% odor concentration (Fig. 2C-D, H). First, we measured the homogeneity of the response signs of the sis-ND6 neurons (Fig. 2E, n=6 animals, 127 glomeruli). When the majority response was an activation, 93.5% of the sis-ND6 also exhibited an activation (t-test, p=0.0034), 89% in the case of an initial inhibition (t-test, p=0.0027; Fig. 2F, n=6 animals, 127 glomeruli). This revealed that sis-ND6 neurons had a higher correlation between themselves than with other cells, an effect which was even more pronounced at lower odor concentration (0.5%, Fig. 2G-I). Therefore, we calculated the average pairwise correlation coefficient of the sis-ND6. We found that it was significantly higher between the sis-ND6 neurons than between the shuffled-neuron groups (Fig. 2J, n=12 animals, 138 glomeruli). Interestingly, the raster plot and pairwise correlation matrix at a lower concentration (0.5%) showed that some sis-ND6 neurons but not all were less well correlated (Fig. 2HI and fig. S4). This pairwise correlation coefficient increased strongly between 0.01% and 20% of the odor concentrations (Chi-squared test, p=0.0164; Fig. 2J).

Using high-resolution in vivo structural imaging, we found that we could efficiently assign the ND6+ neurons to the specific glomerulus onto which they project. Overall, the ND6+ neurons connected to the same glomerulus responded with the same pattern of activity. Surprisingly, we showed that the correlation of sis-ND6 neurons increases with the odor concentration, suggesting a more intricate and nuanced processing of odor presentation. These findings pushed us to further explore intraglomerular representations of odor concentration and identity.

### Intraglomerular dynamic ensembles encode odor concentration and are stable over time

OSNs axons and apical dendrites of MCs in the glomeruli demonstrate correlated responses across a broad range of odor concentrations (*15*). Interestingly, for the sis-ND6 neurons, we observed that their correlation was lower at low concentrations and increased with the concentration range. To further explore this intraglomerular mechanism, we reanalyzed our “odor concentration dataset” using the projection information (n=12 animals, 108 glomeruli with at least 7 sis-ND6 neurons). The sis-ND6 neurons were sorted according to their first responses. We observed that very few cells responded at low concentrations, and the percentage of responses increased with increasing concentrations (Fig. 3A). This progressive recruitment is also striking when we represent the maximum response per cell and per concentration (Fig. 3B).

**Fig. 3.**
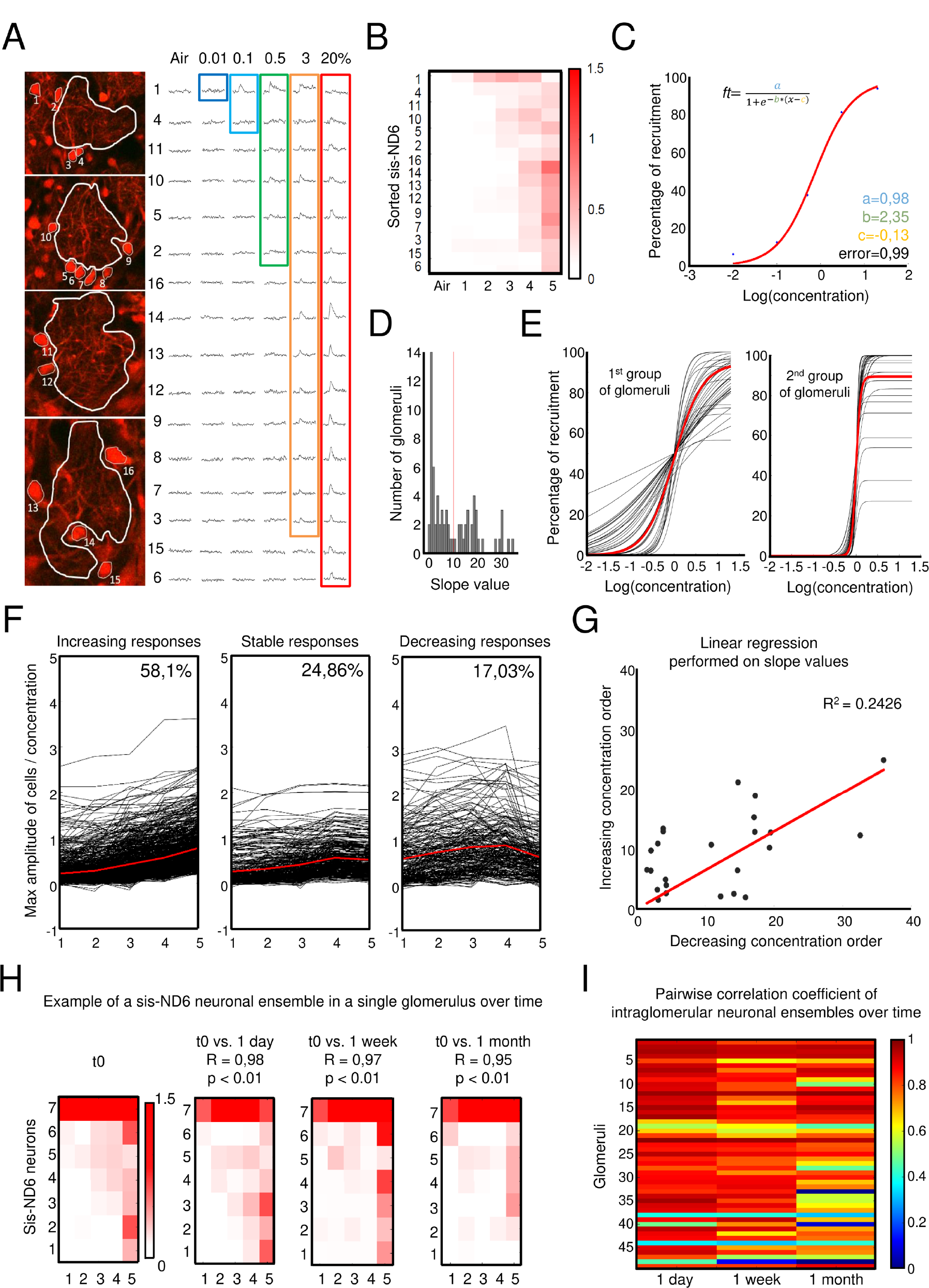
Odor concentration representation by intraglomerular ensembles. (A) Left: Zoomed-in view of a specific intraglomerular ensemble at different planes, with sis-ND6 neurons traced. Right: Calcium responses of sis-ND6 neurons sorted based on their first response within the concentration range. Each block represents sis-ND6 neurons responding to a specific concentration. (B) Raster plot displaying the maximum amplitude of sis-ND6 responses per concentration, as shown in (A). (C) Percentage of cellular recruitment of sis-ND6 neurons, depicted in (A), fitted with a sigmoid model. The mathematical expression of the sigmoid model is provided above. The maximum cell recruitment (a), slope value (b), and EC50 (c) are extracted from the fits. (D) Histogram illustrating the distribution of slope values. The red line indicates the separation in 2 populations by Otsu’s method. (E) Left: Representation of sigmoid fits (normalized by EC50) for neuronal ensembles showing slow recruitment of sis-ND6 neurons. Right: Representation of sigmoid fits (normalized by EC50) for neuronal ensembles showing fast recruitment of sis-ND6 neurons. (F) Plots demonstrating the evolution of ND6 neuron response amplitudes over concentrations, sorted by their monotonicity index value. (G) Linear regression analysis of slope values between the concentration range performed in increasing and decreasing order. (H) Scatter plot showing the correlation of intraglomerular ensembles over time within a specific glomerulus (I) Pseudo-colored representation of pairwise correlation values between intraglomerular ensembles at different time points demonstrating highly stable ensembles (t0 vs 1 day, 1 week, 1 month) (n=5 animals, 49 glomeruli-odor pairs).

As the glomerular response depends on the affinity of the olfactory receptor for the odors presented, in order to compare responses between the glomeruli and between the animals, we fitted the percentage of the recruited cells with a sigmoid model (fig. S5; correct fit for 66% of glomeruli; n=71 over 108 glomeruli). This allowed us to extract a few average characteristics of the responses, independent of the odor affinity (Fig. 3C). The slope of this fit represented the speed of recruitment. We plotted the values of these slopes and distinguished two recruitment profiles (Fig. 3D-E). In 53% of the glomeruli, 28% of sis-ND6 neurons were recruited at 1-log concentration (Fig. 3E, 1st group). In the remaining 47% of the glomeruli, the recruitment was faster, and 49% of the sis-ND6 neurons were recruited at 1-log concentration (Fig. 3E, second group). This indicated that different recruitment speeds exist in different glomeruli. We also quantified the evolution of the response amplitude in individual sis-ND6 neurons across the concentration range. We observed that 58% of the neurons increased their maximum response, whereas 25% remained stable or even decreased their amplitude (17 %; Fig. 3F). To illustrate the stability of this mechanism, we conducted additional experiments involving the repeated presentation of the same odor with a gradual decrease in the concentration (n=5 animals, 39 glomeruli). A linear relationship was observed following the plotting of the slope of the sigmoid fit for both increasing and decreasing concentrations across each glomerulus (Fig. 3G).

To assess the stability of this coding mechanism over time, we recorded the activity of the same neurons at three different time points: 1 day, 1 week, and 1 month later (Fig. 3HI). Then, we applied the same odor concentration range and examined the peak response per cell. We found that the responses remained highly stable over time (Fig. 3H). Both the order of activation and peak response exhibited striking similarities. To quantify this stability, we calculated Pearson’s pairwise correlation coefficient between the glomerular odor peak response maps at each time point. Notably, the correlation coefficient remained consistently high (80% of glomeruli display a significant pairwise correlation coefficient between t0 -1 day, 78% between t0 -1 week, 70% between t0 -1 month), indicating a strong stability of the glomerular responses over time (Fig. 3HI).

Overall, these results showed that odor concentration is encoded by a progressive recruitment of sis-ND6 neurons. Although sis-ND6 neurons connected to the same glomerulus represent a “full” ensemble, only some of them are recruited to adapt to the concentration. We called this mechanism “dynamic intraglomerular neuronal ensembles’’. These dynamic ensembles are not epiphenomena or random activations, but are highly robust and stable over time.

### Dynamic intraglomerular ensembles are also odor-specific

We investigated whether the same intraglomerular ensembles are recruited when the same glomerulus responds to a different odor. To address this, we used two distinct odors and our classical range of concentrations (n=5 animals, 45 glomeruli). Consistent with previous findings regarding the apical dendrites of MCs (*2*), we observed that each odor generated a unique map of active glomeruli (Fig. 4A and fig. S6). Some glomeruli switched from a responsive to a non-responsive state or changed the sign of their response, transitioning from activation to inhibition (Fig. 4A and fig. S6). More importantly, we could already observe on the raster plot that the population and identity of responsive sis-ND6 varied between odors, leading to the emergence of a unique odor response cell map within glomeruli (see. for example Fig. 4A, glomerulus 1). We calculated the pairwise correlations of sis-ND6 neuron responses within individual glomeruli for the two odors, and our findings reveal that these correlations are generally low. Specifically, 69% of the pairwise correlations were found to be non-significant (Fig. 4B, n=5 animals, 45 glomeruli). We focused on the individual glomeruli that responded to the 2 stimulations (30 glomeruli in 5 animals). We sorted the sis-ND6 neurons on the basis of their first response to the first odor, which showed again the presence of concentration-dependent specific neuronal ensembles (Fig. 4C). Strikingly, for the stimulation with a different odor, we observed the activation of a different neuronal ensemble (Fig. 4C-D). We also observed changes in the speed of recruitment of individual glomeruli between the two odors although this was not significant at the population level (Fig. 4E-F, n=5 animals, 14 glomeruli). To quantify these changes, we measured the correlation of the response of sis-ND6 neurons between different odors (Fig. 4G, 2 odors). We show that this correlation was weaker than when compared to the same odor presented on two consecutive days (2 odors compared to t0-t1; p<0.001; Fig. 4G). To assess whether variations in odor affinity might account for the observed differences between the cell recruitment map of the two odors, we shifted the cellular responses to either odor 1 or odor 2 by a factor of 1-log (Fig. 4G, shift odor1, shift odor2). Subsequently, we calculated a new pairwise correlation value based on these adjusted responses. Our results revealed the absence of a significantly different pairwise correlation coefficient. This suggests that the cellular recruitment map we observed was not influenced by odor affinity but rather dependent on the specific identity of the odor itself.

**Fig. 4.**
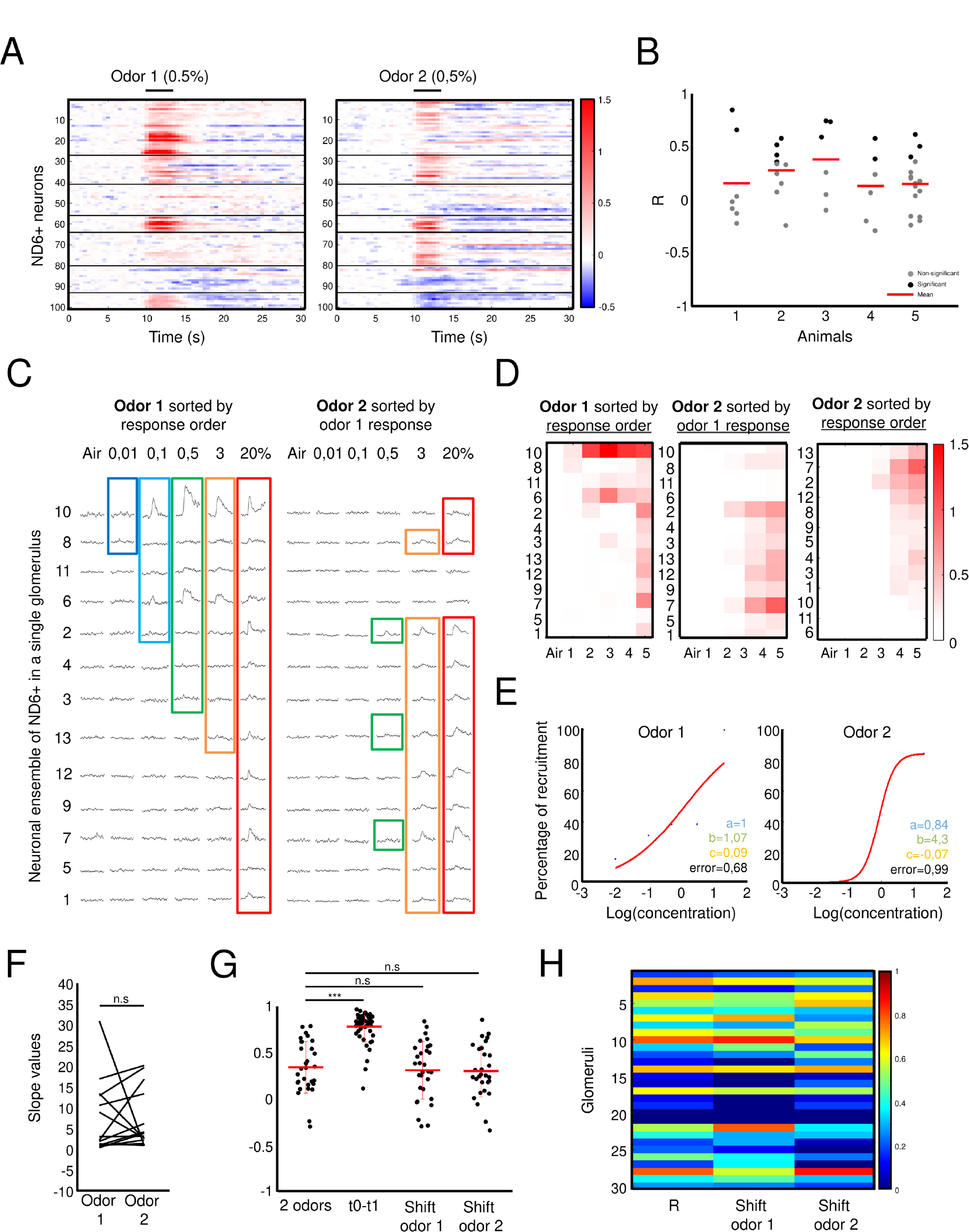
Odor identity is also represented by intraglomerular ensembles. (A) Raster plot illustrating the calcium responses of the same sis-ND6 neurons stimulated with two different odors. The black lines indicate the boundaries of intraglomerular ensembles of sis-ND6. Note the intraglomerular odor specific neuronal recruitment. (B) Intraglomerular cell activity pairwise-correlation between the two odors show that 69% of correlations are non-significant (n=5 animals). (C) Left: Calcium responses of sis-ND6 neurons, sorted based on their first response in the concentration range of odor 1. Right: Calcium responses of sis-ND6 neurons in the concentration range of odor 2, sorted based on the odor 1 concentration range. Note again the highly different recruitment. (D) Raster plot displaying the response amplitude of sis-ND6 neurons per concentration. Left: Response amplitude of sis-ND6 to odor 1, sorted based on response to lowest concentration of odor 1. Middle: Response amplitude of sis-ND6 to odor 2, sorted based on response to lowest concentration of odor 1. Right: Response amplitude of sis-ND6 to odor 2, sorted based on response to lowest concentration of odor 2. (E) Cellular recruitment of sis-ND6 neurons of the same glomerulus for two different odors, as depicted in (A) and (B), fitted with a sigmoid model. (F) Plot showing the slope values for odor 1 and odor 2. The slope is glomerulus and odor dependent. (G) Pairwise-correlation values of intraglomerular ND6+ neurons between two odors. Values between 2 odors are statistically lower compared to the measurement of one odor between two days (Student t-test, ***p<0.001). The pairwise correlation is not different if we shift the concentration range. (H) Same pseudo-colored representation of intraglomerular cell activity pairwise-correlation values between the two odors for 30 glomeruli. ‘Shift Odor1’ indicates the correlation of cell activity maps between the two odors when shifting the concentration range of odor 1 to the right (conc+1). ‘Shift Odor2’ indicates the correlation of cell activity maps between the two odors when shifting the concentration range of odor 2 to the right (conc+1).

Altogether, our results show that odor identity is represented by specific intraglomerular neuronal ensembles.

## Discussion

Our data reveal a new dimension in olfactory information processing, redefining the conventional understanding of odor representation within the OB. To our knowledge, we demonstrate for the first time that odor identity and concentration are represented by a specific subset of glutamatergic neurons in the glomerular layer. We have shown that these neurons orchestrate their responses in a remarkable manner, forming distinct ensembles of sister neurons, all stemming from the same glomerulus. This intraglomerular mechanism could potentially arise from interactions between glomeruli, potentially mediated by inhibitory glomerular interneurons like short-axon cells, as previously proposed (*16*). These interactions may play a crucial role in orchestrating the formation of odor-dependent intraglomerular neuronal ensembles. This, in turn, would facilitate gain control and decorrelation of odor representations, ultimately enhancing the versatility of odor coding within the OB. Overall, we believe that our high-resolution imaging approach, coupled with extensive computational analysis and future modeling, will enable us to gain a better understanding of the intricate processes underlying olfactory information processing.

## Acknowledgments

We thank Michel Picardo, Aurélie Carabalona and Stéphane Bugeon for comments on this manuscript. We also thank the local PiCSL-FBI core facility (Institut de Biologie du Développement de Marseille, Aix-Marseille University) supported by the French National Research Agency through the “Investments for the Future” program (France-BioImaging, ANR-10-INBS-04) as well as the IBDM animal facilities.

## Author contributions

Conceptualization: ML, JCP

Methodology: ML, JCP

Investigation: ML, AA, JCP

Funding acquisition: HC, JCP

Project administration: HC, JCP

Supervision: HC, JCP

Writing – original draft: ML, HC, JCP

## Data and materials availability

The raw data supporting the conclusions of this article will be made available by the authors, without undue reservation.

## Supplementary Materials

Material and Methods

Figs. S1 to S6

Movie S1 to S2

