## supplementary file for "Dynamic intraglomerular neuronal ensembles represent odor identity and concentration"

### Supplementary Materials

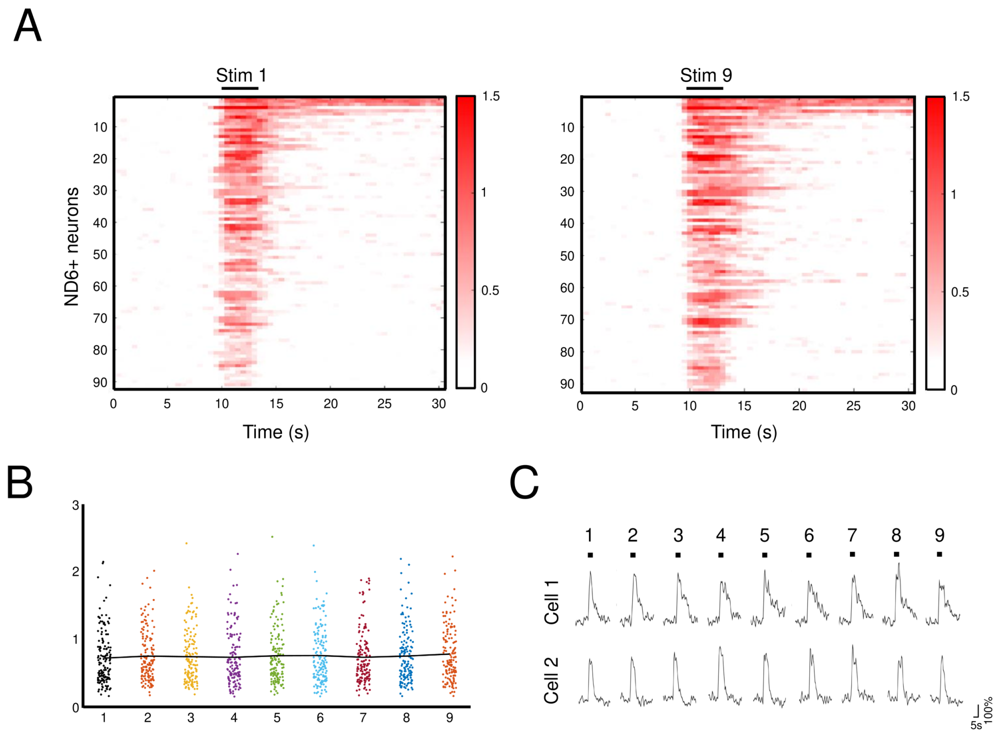


**fig. S1. Passive odor exposure and response dynamics in ND6+ neurons.** (A) Raster Plot illustrating the response dynamics of ND6+ neurons upon repeated stimulation with the same odor (9 times). The raster plots correspond to the initial and final stimulations. (B) Scatterplot presenting the maximum amplitude of the ND6+ neuron responses across multiple stimulations. Statistical analysis indicates non-significant differences between the responses. (C) Representative examples of calcium activity traces observed in two ND6+ neurons upon repeated stimulation with the same odor (9 times).

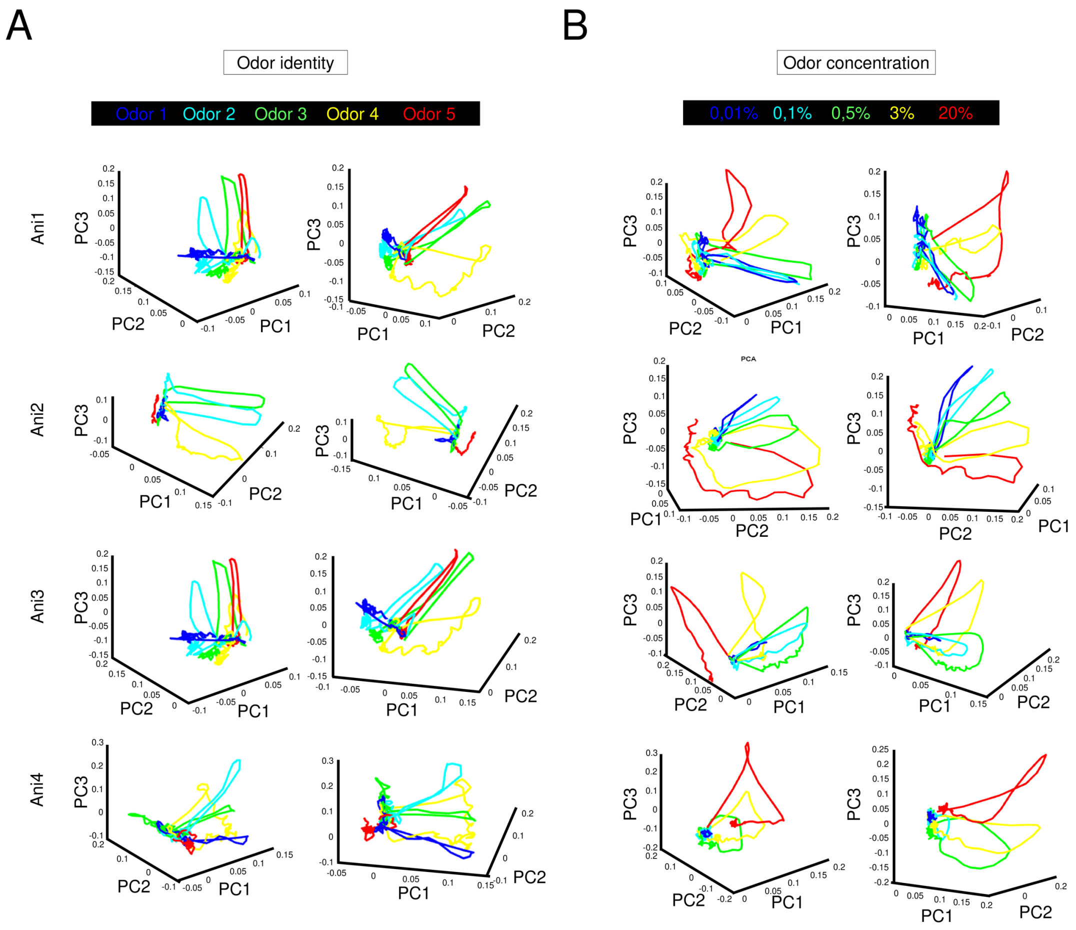


**fig. S2. Principal Component Analysis (PCA) analysis revealing odor identity and concentration coding in ND6+ neurons.** Three-dimensional graphical representations of PC1, PC2 and PC3 for each of the five different odors for 4 animals (A) and 5 concentrations of the same odor (B). Left and right graphs represent two different angles. These graphs show that each odor and each concentration span in a distinct subspace, indicating that odor identity and concentration are encoded in this population in the glomerular layer.

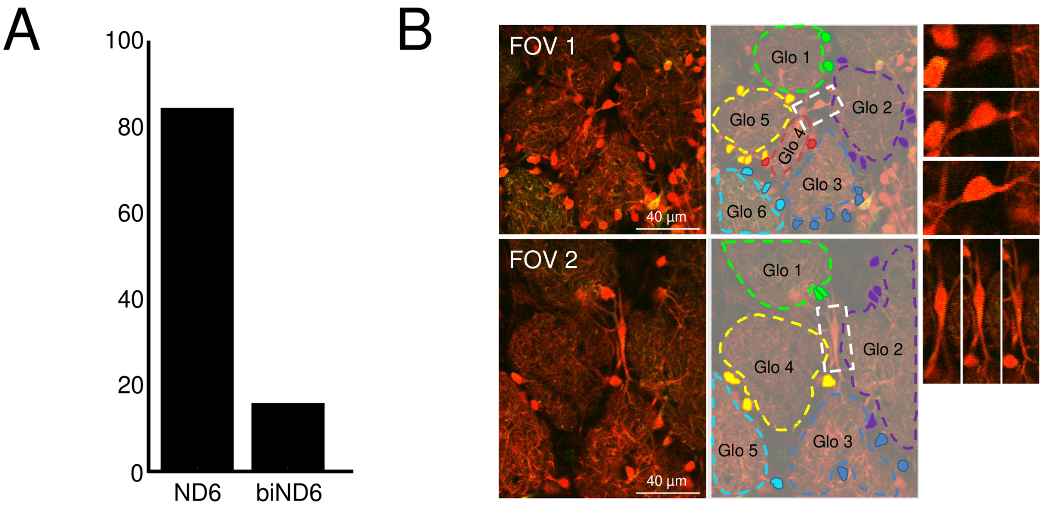


**fig. S3. Morphologies of ND6+ neurons in the glomerular layer.** (A) Bar graph depicting the percentage of ND6+ neurons based on the projection pattern of their main dendrite showing that most ND6+ neurons have only one main dendrite projecting within a single glomerulus (n=5 animals). (B) Two examples of ND6+ neurons with their main dendrite projecting in two glomeruli.

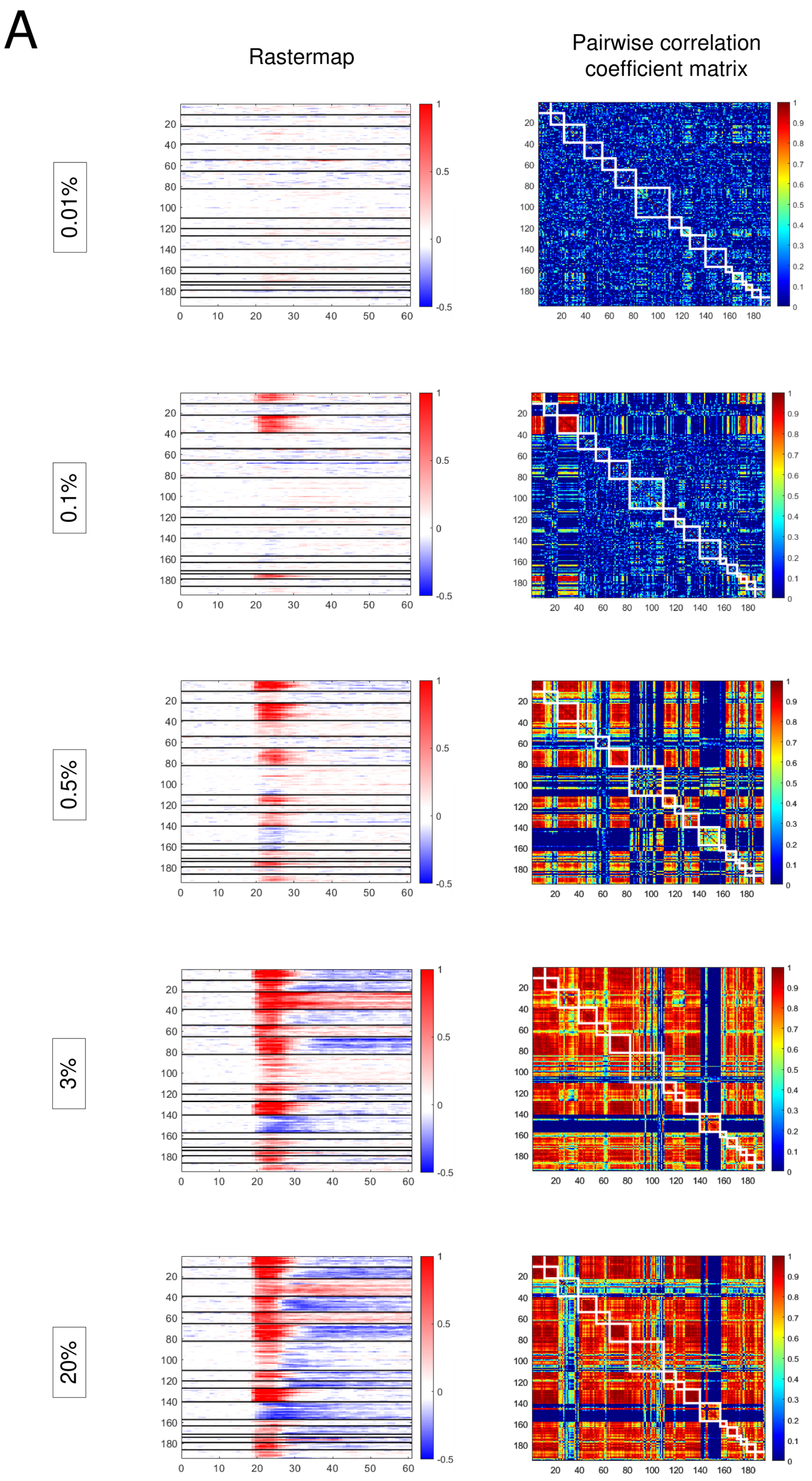


**fig. S4. Evolution of intraglomerular neuronal ensembles response through a concentration range.** (A) Glomerular specific recruitment as a function of odor concentrations (0.01% to 20%) (B) Pearson pairwise-correlation matrices illustrating the correlation of sis-ND6 responses and how it evolves with odor concentration.

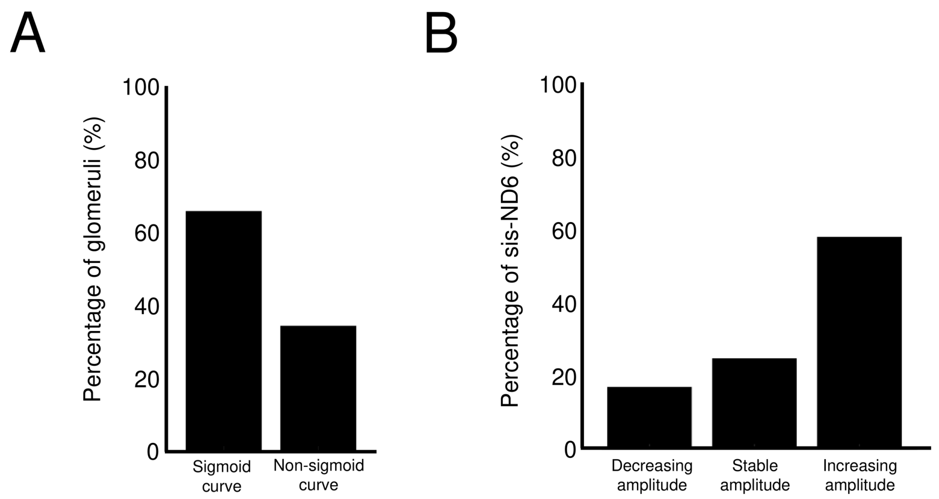


**fig. S5. Behavior of sis-ND6 neurons across a concentration range.** (A) Bar graph representing the percentage of glomeruli where cell recruitment fits accurately using a sigmoid function. (B) Average evolution of response amplitudes observed within intraglomerular neuronal ensembles.

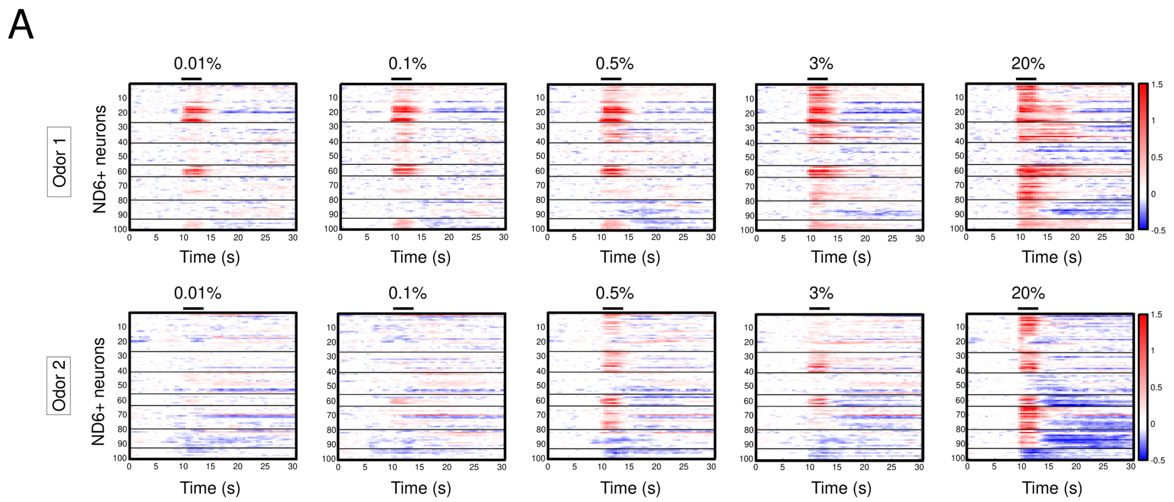


**fig. S6. Evolution of responses within intraglomerular neuronal ensembles of the same animal between two different odors across the same concentration range.** The raster plot visually represents the temporal dynamics of responses within intraglomerular neuronal ensembles as observed between two different odors. The concentration range remains constant for both odors.

**Movie. S1. High-resolution Z-stack of the glomerular layer performed in an ND6-Cre/tdTomato mouse model.** Images were acquired using a field of view of 600m² with a 2 µm step size (resolution of 1024x1024 pixels).

**Movie. S2. High-resolution Z-stack of sis-ND6 within a unique glomerulus.**

### Material and Methods

#### Animals

All mice were treated according to protocols approved by the French Ethical Committee (#5223–2016042717181477 v2). Mice were group housed in regular cages under standard conditions, with up to five mice per cage on a 12 hr light–dark cycle. NeuroD6Cre (ND6Cre) knock-in mice (Goebbels et al., 2006) were bred with Rosa26tdTomato reporter mice (Ai14, Jackson Laboratories, stock-number 007914) and GCaMP6s mice (Jackson Laboratories, stock-number 007914). Mice of both sexes were used.

#### Implantation of cranial window

Implantation of the observation window was performed as previously described (Drew et al., 2010) with some modifications. Mice were anesthetized by intraperitoneal injection of ketamine/xylazine (125/12,5 mg/kg). Dexamethasone (0.2 mg/kg) and buprenorphine (0.3 mg/mL) were injected subcutaneously 30 minutes before the surgery and lidocaine was applied locally onto the skull. The pinch withdrawal reflex was monitored throughout the surgery, and additional anesthesia was applied if needed. Carprofen (5 mg/kg) was injected after the surgery. A steel bar was added during this step to allow fixation of the animal to the microscope. The skull overlying the OB was carefully thinned with a sterile scalpel blade until a thickness of 10–20 µm was reached. A thin layer of silicon was applied on the window and a 3-mm round coverslip was apposed and sealed with glue (Vetbond) and dental cement (superbond, GACD). A first microscopic observation was performed on these anesthetized mice.

#### In vivo two-photon imaging

We used a Zeiss LSM 7MP two-photon microscope modified to allow animal positioning under a 20X water immersion objective (1.0 NA, 1.7 mm wd) and coupled to a femtosecond pulsed infrared tunable laser (Mai-Tai, SpectraPhysics). Epifluorescence signals were collected and separated by dichroic mirrors and filters on four independent non-descanned detectors (NDD). Images were acquired using an excitation wavelength of 970 nm. In general, image acquisition lasted about 1 hour under ketamine/xylazine (125/12,5 mg/kg) anesthesia. The imaging window was centered on the dorsal surface of the OB. In cases where consecutive observations were necessary for the same animal, the same field of view was identified based on the arrangement of glomeruli and groups of sis-ND6, as well as specific morphological features of individual cells. Before initiating the odorant stimulation protocol, a Z-stack of the entire field of view (approximately 250 µm depth) was acquired using a step size of 2 µm in the Z dimension and a xy pixel size of 0.6 µm. This approach allowed us to identify the focal plane for calcium imaging within the Z-stack, accurately reconstruct and assign intraglomerular cells to their respective glomeruli. Calcium imaging was conducted at three distinct dorsoventral levels within the glomerular layer, with a separation of 20 µm between each level using 512x512 resolution (frame rate of 2 Hz). This sampling strategy allowed for comprehensive assessment of sis-ND6 neurons activity within the glomeruli based on odor identity and concentration. The reconstructed sis-ND6 data were processed using FIJI software (Schindelin et al., 2012), while calcium transients were analyzed using a custom-made Matlab analysis pipeline.

#### Olfactometry

Odors were delivered using a 6-valve liquid dilution olfactometer (Knosys Olfactometers). Commercially available odorant molecules were diluted in mineral oil to obtain concentrations between 0.01% and 20%. Odors used in this study are known to activate dorsal glomeruli at these concentrations (Adam et al., 2014) : Butanal, Ethyl Tiglate, Methyl Propionate, Propanal, Butanal, Pentanal, Hexanal, Ethyl Tiglate, Ethyl Butyrate, Methyl Benzoate, Methyl Propionate, Hexyl Acetate, Isoamyl Acetate, Isoamyl Butyrate, Carvone. Olfactometry: All odors used in this study were obtained from Sigma-Aldrich with a purity level of >98%. The experimental protocol involved a sequence of events: an 8-second delivery of exhaust, followed by a 3-second odor stimulation, and finally, 40 seconds of interstimulation-intervals (ISI). To ensure accurate odor delivery, the odor delivery device was positioned approximately 1 cm in front of the animal's nose. An auxiliary constant air flow system was placed on the side to effectively remove any residual odor after each presentation. Prior to each session, animals were exposed to clean air as a control stimulus. Rare cases of glomeruli that responded to clean-air stimulation were not analyzed.

#### Histology

For histological analysis, mice were deeply anesthetized with an xylazine/ketamine overdose. Intracardiac perfusion was performed with 4% paraformaldehyde in PBS. The brain was collected and incubated overnight in the same fixative at 4°C. Sections were cut at 50 µm sections were cut with a microtome (Microm). Standard immunostaining protocols were performed on coronal free floating sections. Briefly, sections were rinsed in PBS and incubated in a blocking buffer [10% fetal bovine serum (FBS), 0.3% Triton X-100 in PBS] for 1 h. Subsequently, sections were incubated in primary antibody solution [5% FBS, 0.1% Triton X-100 in PBS (PBST)] and VGLUT2 antibody [Synaptic system, 1:2,000] overnight at 4°C. The following day, sections were rinsed 3 times in PBS and incubated with species-appropriate secondary antibody in PBST for 2 h at room temperature using gentle rocking. Alexa Fluor conjugated secondary antibodies were from Jackson ImmunoResearch. Nuclear counterstain HOECHST 33258 (Invitrogen, 1:2,000) was added before sections were washed in PBS and mounted using Mowiol as a mounting medium. Optical images were taken with a laser confocal scanning microscope (LSM780, Zeiss, Germany) and image analysis was performed using FIJI software (Schindelin et al., 2012).

#### Calcium activity trace processing and odor response detection

The calcium activity traces were extracted from manually drawn masks using Fiji. The traces were first normalized by dividing the fluorescence intensity of the green calcium indicator (GCaMP6s) by the fluorescence intensity of the red indicator (tdTomato, considered stable in the cells) to account for motion artifacts. A running average of 2 images was then applied to denoise the signal. The resulting ratio traces were analyzed for significant responses to olfactory stimulation.

To determine the response threshold, the maximum response amplitudes were calculated for each region of interest (ROI) during the 3 seconds stimulation. Excitation was detected if 1.the maximum amplitude exceeded the maximum response amplitude during air stimulation 2. was higher than three times the standard deviation (SD) in the pre-air stimulation period and pre-olfactory stimulation period, and 3. had a value higher than 0.15. Inhibitions were described to have lower amplitude so we used a different threshold. They are detected if 1. the absolute value of the minimum response amplitude exceeds the maximum response amplitude during air stimulation, 2. were higher than two times the SD at the air pre-stimulation period and olfactory pre-stimulation period, and 3. had a value higher than 0.15. A response was classified as complex if both excitation and inhibition criteria were met.

#### Passive odor stimulation

We investigated the sensitivity of ND6 positive neurons to passive odor stimulation on anesthetized animals using the same focal plane and multiple stimulations at a concentration of 1%. This odor stimulation was repeated nine times consecutively. The protocol was replicated for different odors, allowing for the recording of a maximum number of responding neurons during a single session. For the analysis, we focused only on cells that exhibited a response to olfactory stimulation during the first trial. This criterion ensured that we captured and analyzed cells that were specifically responsive to the odor stimulus. To evaluate the statistical significance, a repeated measures ANOVA was conducted using the ‘ranova’ function.

#### Principal Component Analysis of odor identity and concentration

PCA was performed using MATLAB on odor identity or concentration dataset. The analysis focused on the response of the global population of glomerular ND6 to odor stimulation. We used the whole stimulation epoch (10 seconds pre-stimulation, 3 seconds stimulation and 42 seconds post-stimulation). We denoised the signal using a Savitzky-Golay filter with a length of 5 frames (2.5s). PCA was then performed on the preprocessed intensity data using the "pca" function in MATLAB. We plotted in 3 dimensions the first three principal components. Each odor or concentration level was represented by a distinct color. The plot displayed the trajectories of the odor or concentration responses over time, allowing for a visual assessment of the similarities and differences between different odor conditions or concentrations.

#### Non-Negative Matrix Factorization analysis of Odor concentration

To investigate the odor concentration dynamics in the ND6 population, a Non-Negative Matrix Factorization (NNMF) analysis was conducted using the MATLAB “nnmf” function. To evaluate the quality of the NNMF decomposition, the approximation error (using the Frobenius norm) was calculated for a range of ranks (1 to 20). The spatial distribution of the NNMF components was examined. The glomerular structure image was overlaid with the glomerular

ROI displayed in white. Each NNMF component was analyzed to identify regions with weights above a specified threshold. The corresponding ROIs were plotted on the glomerular image in red, providing insight into the spatial distribution associated with each NNMF component.

**Comparison of odor response in sis-ND6**

To quantify the similarity of intraglomerular cell responses, we calculated the Pearson pairwise-correlation coefficient. We specifically focused on glomeruli that exhibited neuropil responses to olfactory stimulation between 0.01 and 20 % (n= 12 animals, 171 glomeruli, 2485 neurons). For each animal and each responsive glomerulus, we calculated the correlation coefficient based on the response traces of all sis-ND6 (Fig. 2J). This coefficient represents the degree of linear relationship between the responses of these cells. To establish the statistical significance of the observed correlation values, we employed a permutation test. We conducted 10 000 simulations, wherein we randomly selected cells that did not project into the same glomerulus. Subsequently, we recalculated a new correlation value based on the response traces of these shuffled cells for each simulation (as depicted in Fig. 2J). The p-value was then calculated as the proportion of simulations in which the "shuffle" value exceeded the "real" value divided by the total number of simulations (10 000). This provided a measure of the statistical significance of the observed correlation between intraglomerular cell responses, indicating the likelihood of obtaining such a correlation by random chance.

**Comparison of intraglomerular cell activity map over time**

To evaluate the stability of intraglomerular activity maps over time, the Pearson correlation coefficient is computed based on the maximum amplitudes of olfactory responses in sis-ND6 obtained at different time points (Fig. 3HI). Specifically, the Pearson coefficient is measured between the responses obtained at time t0 and the subsequent time points (1 day, 1 week, and 1 month later), using the maximum amplitudes of the responses.

**Comparison of intraglomerular cell activity map between odors**

To assess the similarity of intraglomerular cell activity maps between two different odors, first, we calculated the Pearson pairwise-correlation coefficient. To check whether variations in odor affinity might account for the observed differences between the cell recruitment map of the two odors, we shifted the cellular responses to either odor 1 or odor 2 by a factor of 1-log. Subsequently, we calculated a new pairwise correlation value based on these adjusted responses.

**Fitting of odor responses**

A sigmoid model was used to fit the cellular recruitment data within glomeruli. Experimental data consisting of log-transformed concentrations (logconc) and corresponding cellular recruitment values (x) were collected. The sigmoid model equation used in this study is as follows:

$$x = a / (1 + \exp(-b * (\text{logconc} - c)))$$

'a' represents the maximum recruitment, 'b' represents the slope, and 'c' represents the EC50 value. The EC50 value corresponds to the logconc value at which the cellular recruitment reaches half of its maximum value.

The sigmoid model was fitted to the experimental data using a nonlinear least squares optimization method. After the model fitting process, a fit object and a goodness-of-fit structure were generated. The coefficient of determination (R-squared) was calculated and if less than 0.6, it is considered that the sigmoid function is not a suitable model, indicating the absence of progressive cell recruitment within the glomerulus.

To better quantify the cell recruitment mechanism within glomeruli, we employed Otsu's method to determine an optimal threshold for classifying them into two distinct populations based on their slope values. This approach allowed us to effectively distinguish and categorize glomeruli into their respective recruitment speed groups, enabling a more accurate analysis of the cell recruitment dynamics within the studied samples.

**Monotonicity index**

In this analysis, we focused on cells that exhibited responses at some point across the concentration range in order to analyze the evolution of their maximum response amplitudes (Fig. 3F). As our experimental design involved olfactory stimulations with increasing concentrations, we expected the maximum response amplitude of each cell to occur at the highest concentration. To determine this, we employed an index score calculation based on the method described by Escabi et al. (2007):

D = (Maximum amplitude at concentration Cx - Maximum amplitude at highest concentration Cmax) / standard deviation (sd)

By analyzing the value of D, we could determine the concentration at which the cell reached its maximum amplitude. If D = 0 was reached at the last concentration, this suggests a direct and proportional relationship between the stimulus and the cell's activity.

However, if D = 0 was reached before the last concentration, we made further considerations. If the minimum absolute value of D after reaching zero was lower than -2 times the standard deviation ($|D| < -2sd$), we concluded that the response amplitude of the cell maintained a stable level from that particular concentration onward. Conversely, if the minimum absolute value of D after reaching zero was higher than -2 times the standard deviation ($|D| > -2sd$), we interpreted it as a decrease in the response amplitude of the cell from that concentration onward.

**Statistics**

Statistical comparisons were performed using Matlab software (Mathworks). Threshold for significance was set at p=0.05 (*p<0,05, **p<0,01, ***p<0,001). To assess the differences in the number of responding neurons across different concentrations, we employed a Cochran-Armitage trend test (***p<0.001). We assessed the homogeneity of sister-ND6 odor responses within glomeruli (Fig. 2E) using a shuffling method with 10 000 simulations per glomerulus. The p-value for each glomerulus was determined based on the number of times the shuffle value exceeded the correlation value of Pearson's coefficient. In Fig. 2F, we additionally used a t-test that measured the significance of values of Pearson’s coefficient with the mean value of shuffling values (10 000 simulations) : p=0.0034 for activations, p=0.0027 for inhibitions. To evaluate the correlation of odor responses across different odor concentrations (Fig. 2J), we compared Pearson's coefficient value with shuffling values generated through 10 000 simulations. The p-value for each glomerulus was calculated according to the number of times the shuffle value exceeded Pearson's coefficient value. Additionally, we employed a Chi-squared test to examine the evolution of the distribution of significant glomeruli across odor concentrations ($p=0.0164$). We performed linear regression analysis to compare the values of cell recruitment slope between increasing and decreasing odor concentrations (Fig. 3G). The significance of Pearson’s coefficient values, both over time (Fig. 3HI) and between different odors (Fig. 4B), was assessed using the method described in Fig. 2EJ. To compare slope values between different odors (Fig. 4F), we employed a t-test, yielding a non-significant p-value of 0.8736. To compare the Pearson’s coefficient values between odors and time (Fig. 4G), we employed a student t-test (***$p<0.001$). To compare the Pearson’s coefficient values between odors and ‘Odor shift’ (Fig. 4G), we used a t-test, yielding a non-significant p-value of 0.6775 (odors - Odor 1 shift) and 0.5745 (odors - Odor 2 shift).
